# Improving Lab Culture through Self-Assessment: A Case Study

**DOI:** 10.1101/2021.12.08.471870

**Authors:** Soleil Hernandez, Raymond Mumme, Laurence Court, Daniel El Basha, Skylar Gay, Barbara Marquez, Yao Xiao, Kai Huang, Hana Baroudi, Wenhua Cao, Carlos Cardenas, Raphael Douglas, Jack Duryea, Zaphanlene Kaffey, Deborah Mann, Kelly Nealon, Tucker Netheron, Callistus Nguyen, Kyuhak Oh, Adenike Olanrewaju, Carlos Sjogreen, DJ Rhee, Jinzhong Yang, Cenji Yu, Lifei Zhang, Yao Zhao, Hamid Ziyaee, Mary Gronberg

## Abstract

**Purpose:** Motivated by perceived dissatisfaction within our lab’s changed working environment brought about by the COVID-19 pandemic, we performed a self-assessment of our lab culture through anonymous surveys and live sessions.

**Methods:** In Survey 1, we asked each lab member to identify and rank up to 10 values that are important for a healthy lab environment. They were then asked to rate how well the lab embodied those values at two time points: before the COVID-19 pandemic while working onsite, and at the time of the survey while working remotely (10 months into the pandemic). In a series of live group sessions, we reviewed relevant literature and the survey results to finalize ten themes. We then reflected on each theme and proposed action items to address any deficiencies. Finally, we conducted Survey 2 after the self-assessment to judge the group’s finalized themes, implemented changes, and overall satisfaction with the assessment process.

**Results:** Themes identified were attitude, accountability, teamwork/collaboration, communication, diversity/inclusion, emotional intelligence, integrity, training, well-being, and adaptability in crisis-management. All lab members liked the self-assessment process and felt their voices were heard. On average, there was a 12% increase in satisfaction across all themes from the start to end of the lab assessment.

**Conclusion:** We successfully assessed the culture of our lab and subsequently improved lab member satisfaction. The success of this team project suggests that other scientific labs could benefit from similar interactive self-assessments.

## Introduction

A healthy work environment has been shown to increase overall team morale and subsequently reduce turnover rates(1) and improve productivity and creativity(2,3). Similarly, a healthy scientific lab environment fosters good science(4,5) and improves collaboration(6,7). Unfortunately, several surveys reveal the prevalence of unhealthy lab environments. For example, a recent Nature survey of 3,200 scientists highlights discrepancies in presumed workplace satisfaction between lab leaders and reported workplace satisfaction by lab members(8). Another Nature survey of over 6,300 early-career researchers revealed frustration with quality of training, work-life balance and cloudy job prospects(9).

Previous studies have recommended strategies to effectively mentor trainees, foster productivity, and overcome unexpected challenges(10,11). In addition to recommending strategies, several research labs have published “rules” for a healthy lab environment(12–14). Examples of these rules include promoting collaboration, gratitude, work-life balance, and professional development. While these rules are applicable to most lab environments across disciplines, each lab has a unique culture and prioritizes different values(15,16). To further promote lab satisfaction, it is important that lab values reflect the lab members’ beliefs and not just the experience of the lab manager(8). With this in mind, we performed a self-assessment of lab culture, including identification of lab values, evaluation of how well we embody our lab values, development and implementation of necessary changes, and recommendations for future continual assessment(14). In this work, we detail our lab’s self-assessment as a case study to provide a translatable framework for evaluating and improving lab culture. We describe the results of the assessment, including the themes identified, and the evident, positive effect on our lab’s environment.

## Materials and Methods

An overview of our self-assessment approach is detailed in Figure 1. To self-assess our lab culture, we began with a survey to start to identify values of the lab members. We then discussed the results of the survey over a series of interactive live sessions. In these live sessions, we identified weak points within our lab and proactively proposed and enacted solutions to address them. Finally, we conducted a second survey to determine the impact of this process.

**Fig 1.**
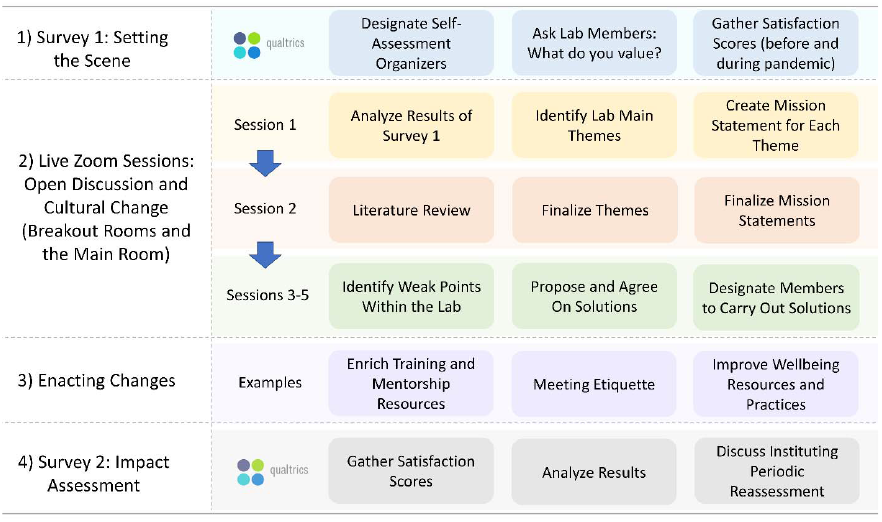
Overview of our methodology for assessing our lab culture.

IRB approval was not obtained for this study because we do not consider this work to be human subject research as all survey participants were members of our lab and authored this work.

### Survey 1: Setting the Scene

An initial survey was conducted anonymously using Qualtrics, in which lab members were each asked to identify and subsequently rank a minimum of 3 and a maximum of 10 values that are important when searching for a healthy lab environment. Respondents were then asked to score how well the lab environment embodied their contributed values at two time points: prior to the COVID-19 outbreak while working onsite, and at the time of survey while working remotely (10 months into the COVID-19 pandemic). This satisfaction score was evaluated on a scale of 1-5, with 5 indicating the best manifestation of the value within the lab’s culture. The full survey can be found in the S1 Appendix.

### Sessions 1 & 2: Open discussion

During Session 1, our lab group met virtually over Zoom for three hours to discuss the results from Survey 1 and to develop themes that encapsulated all lab members’ values. This Session was divided into three portions: a comprehensive review of the survey results, individual breakout room sessions to discuss the results, and combined group sessions to compile ideas and refine our themes.

The results of Survey 1 were compiled and visually presented to all lab members. Team members were divided into breakout sessions in Zoom to sort values into overarching themes. After each group session, we reconvened in the main Zoom session. To organize our findings and ideas, each group provided a summary of their discussions, including voiced ideas and potential amendments to the current list and definitions of proposed themes. These larger discussions led to the establishment of our initial themes and their corresponding mission statements.

The objectives of Session 2 were to perform a lab culture literature review and use our findings to finalize our lab’s themes. It should be noted that the literature review was performed after Session 1 to avoid biased Survey 1 responses. To perform the group literature review, lab members were divided into four work groups, each tasked with presenting at least two lab culture articles and a perspective article(8,10,22–25,12– 14,17–21). From our literature review, we refined our set of themes. For each theme, we finalized their mission statements and identified associated key values.

### Sessions 3-5: Cultural Change

The objectives of Sessions 3-5 were to identify weak points within the lab and propose solutions. To accomplish this, we followed these steps:

1. For each theme, the aggregated Survey 1 results were summarized and presented for reference.
2. Using polling software (PollEverywhere), we asked participants to answer the following questions for each theme: (1) “What do we do wellã” and (2) “How can we improveã”. The software allowed the process to be anonymous and allowed participants to agree or disagree with others’ responses in real time.
3. We reflected on the responses of the poll and identified key deficiencies. We then split into small groups to brainstorm new ideas to address these deficiencies.
4. After small group sessions, we met as a large group to record the action items to improve lab culture. Specific action items were assigned to individuals to implement the changes as appropriate.

### Survey 2: Impact Assessment

A second survey was conducted three months after the last live session to assess the finalized lab themes, the implementation outcomes, and the entire self-assessment process. Lab members were also asked (at the time of this second survey, while still working remotely) to rate how well the lab embodies their values from the first survey using the 1-5 scale. This satisfaction score at a third time point was collected for comparison to the satisfaction scores at the other two time points from the first survey. All survey questions can be found in the S1 Appendix.

## Results

### Sessions 1 & 2: Open discussion

In Sessions 1 and 2, we finalized our central themes and mission statements. After analysis of the results from Survey 1, the 134 submitted values by 21 lab members were sorted into primary themes. Following group discussion, the primary themes were renamed or combined into finalized themes. Adaptability in crisis management was an addition to the finalized themes resulting from the literature review. As a result, we finalized 10 central themes for our lab.

The process of identifying our final themes is detailed in Figure 2. We separated into small breakout rooms (4-5 members each) to craft mission statements for each theme. The mission statements and corresponding values are listed for each theme in Table 1.

**Table 1.**
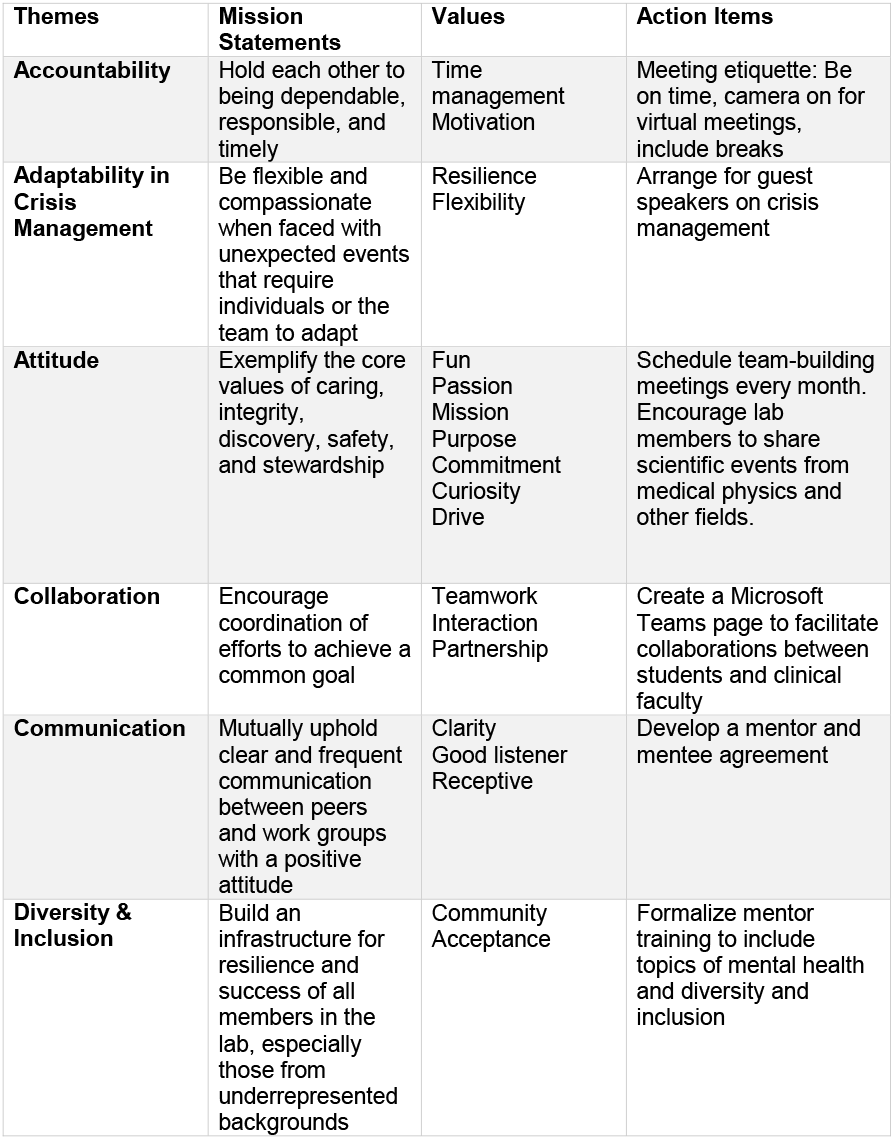
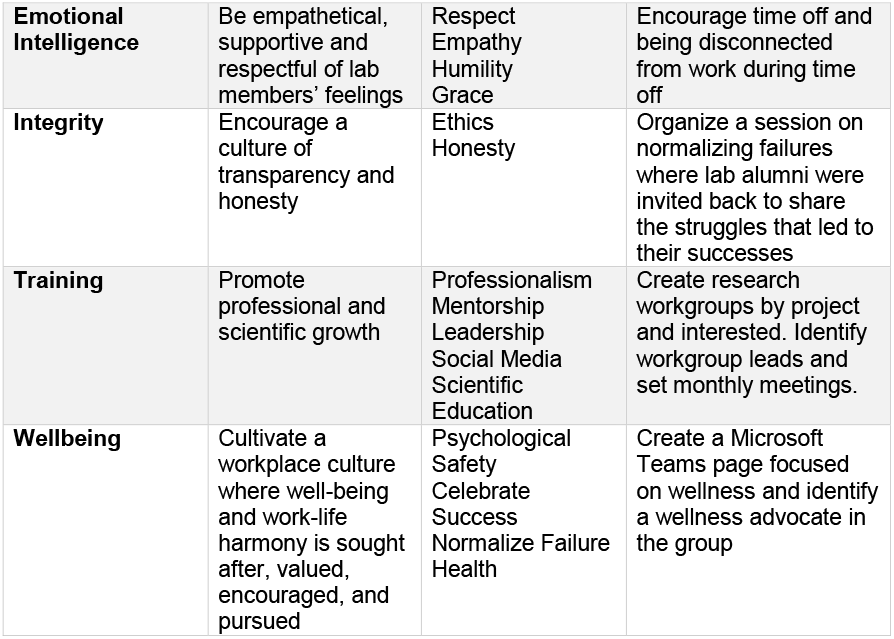
Lab themes and their corresponding mission statements, values, and proposed action items.

**Fig 2.**
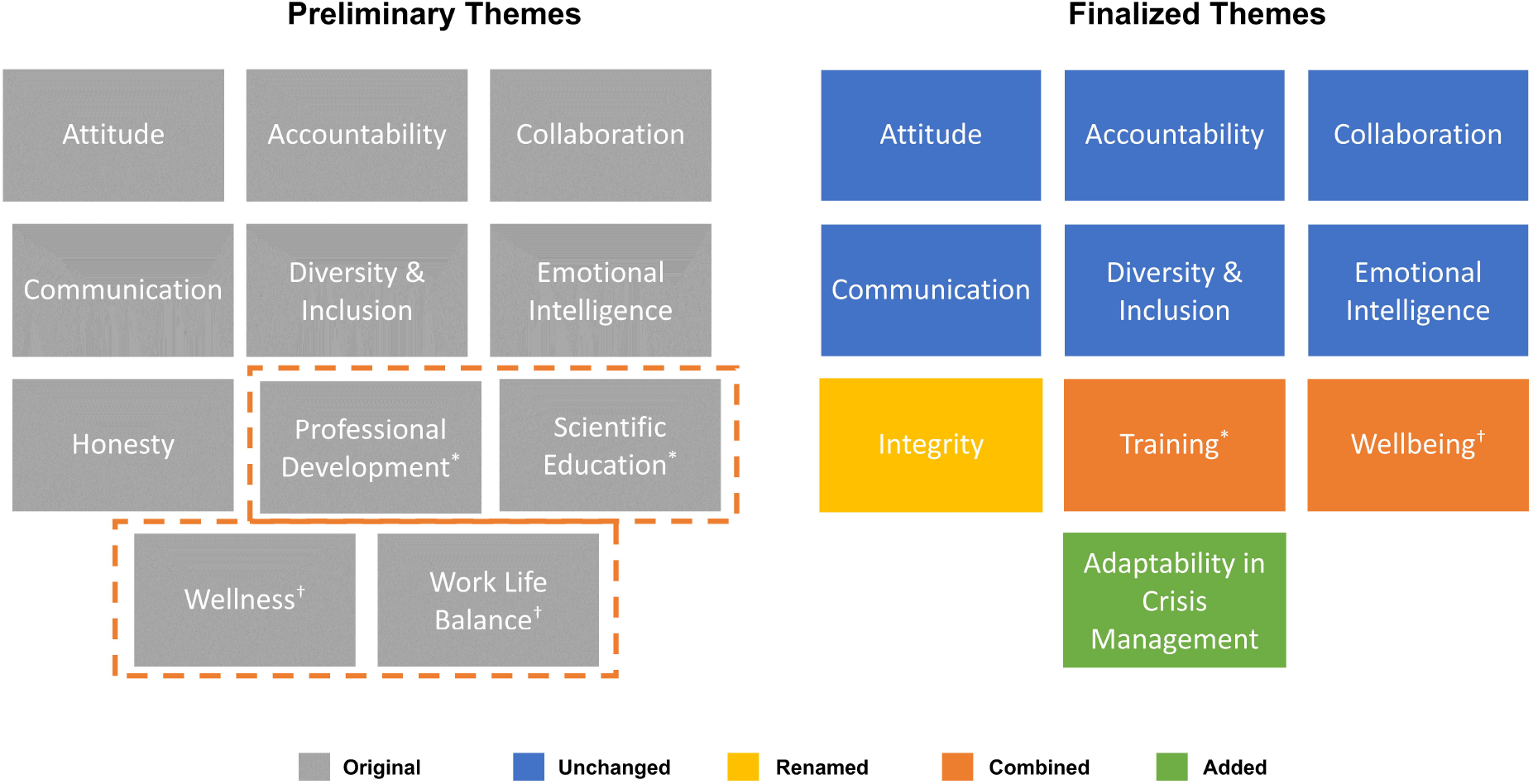
Comparison of the primary themes that emerged from Survey 1 and the finalized themes that resulted from live discussion.

### Sessions 3-5: Cultural change

In Sessions 3-5, we identified weak points within the lab culture and proposed solutions to address them. The discussions resulted in at least one action item per theme. Example action items are shown in Table 1. Following the live sessions, specific team members were assigned to follow through with these action items.

The two themes that were identified as the weakest points in our group were accountability and wellbeing. For accountability, students felt that transitioning to a remote working environment (because of institutional COVID-19 precautions) made it more difficult to stay motivated and manage their time wisely. To address this issue, several action items were proposed by each small group. First was meeting etiquette. Members noted the importance of being on time for meetings and keeping meetings brief to maximize attention spans. We implemented ‘read receipts’ on important lab announcements and emails by having each member acknowledge that they had read an important post. Finally, we created open Zoom rooms for virtual work to simulate working alongside coworkers in the office. For wellbeing, students reported that they had difficulties establishing boundaries between work and their personal lives. To address this issue, the students proposed creating a wellness channel on teams and identifying a wellness advocate in the group. The wellness channel allowed for each member to post and view various wellness resources including exercise, meditation, mental health, cooking, and other hobbies. Dedicating a team’s channel to the lab members’ personal lives helped cultivate community within the group. Additionally, we created a trainee group chat on teams to encourage trainee interaction beyond technical questions.

### Surveys 1 & 2

The 21 lab members who participated in Survey 1 also participated in Survey 2 to provide their input on the lab culture assessment. Participants ranked the 10 finalized lab themes in order of importance to them. The rankings are displayed in Figure 3. The top ranked themes across the lab were collaboration, accountability, and training. Accountability and training were ranked 1 most frequently, followed by integrity. Wide distributions in rank were observed (Figure 3), emphasizing that every lab member prioritizes values differently and that efforts to maintain lab culture should be balanced across all themes.

**Fig 3.**
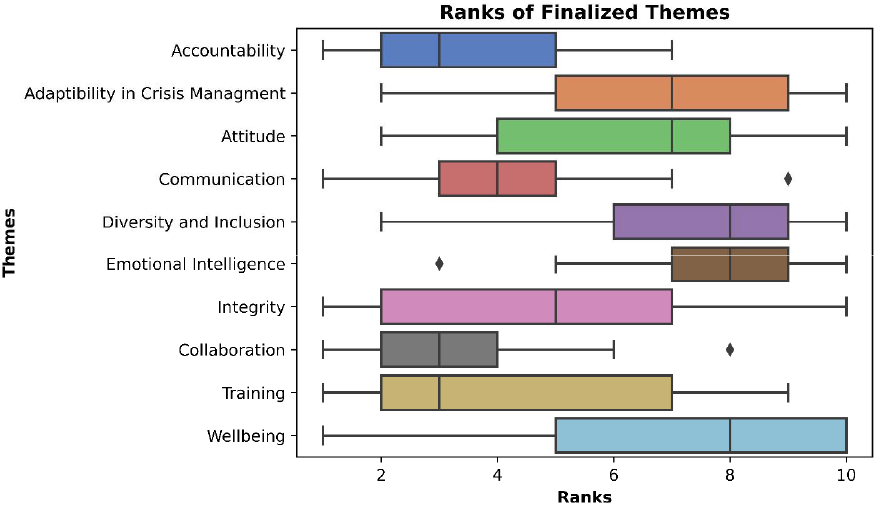
Distribution of lab members’ ranking for each finalized theme.

Figure 4 shows changes in average lab satisfaction scores across identified themes at three time points: (time point 1, blue) before the pandemic while working onsite; (time point 2, orange) 10 months into the pandemic while working remotely, prior to the lab self-assessment; (time point 3, green) and 5 months after the second time point while working remotely, at the conclusion of the lab assessment. Because adaptability in crisis management was added after time point 2, it is not displayed in Figure 4. The adaptability in crisis management rating at time point 3 was 4.48 ± 0.51.

**Fig 4.**
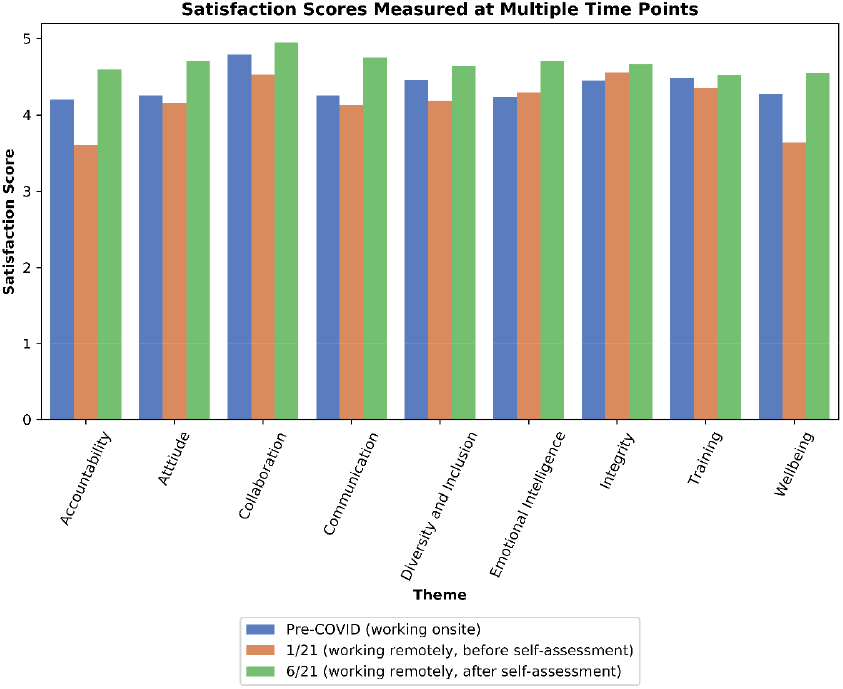
Ratings of original values grouped by themes and measured at three time points. Increased satisfaction scores were reported across all finalized themes

The average satisfaction scores averaged over all themes at time points 1, 2, and 3 were 4.37 ± 0.78, 4.16 ± 0.93, and 4.67 ± 0.51, respectively. The themes with the largest decreases in satisfaction scores between time points 1 and 2 were accountability and wellbeing (14% and 15%, respectively). There were increases in satisfaction scores for accountability and wellbeing between time points 2 and 3 following the lab culture assessment (28% and 25%, respectively). On average, there was a 12% increase in satisfaction across all themes from the start to end of the lab assessment (between the second and third time points). Additionally, we saw an overall average 7% increase in satisfaction across all themes from prior to the pandemic when our lab was working onsite to after the self-assessment (between the first and third time points).

All participants reported feeling that their voice was heard throughout the process, with 57.1% agreeing to a high degree. Also, nearly half (47.6%) of the survey participants thought that any changes made after Survey 1 helped them improve their working conditions. All participants believed that these changes would be sustained in the long term if we reviewed our lab culture periodically.

A majority (57.1%) of respondents liked the assessment of the lab culture to a high degree, with an additional 33.3% and 9.5% liking it to a lesser degree or only somewhat liking it, respectively, while no lab members reported disliking the assessment. Similar results were observed for the reasonableness of the time commitment required by this assessment, with 85.7% and 9.5% feeling it was reasonable or somewhat reasonable.

## Discussion

In this study, we demonstrated that an interactive self-assessment of lab culture can give an overall increase in satisfaction across core lab values (categorized into themes). Conducting this self-evaluation was motivated by perceived dissatisfaction within the lab’s changed working environment (i.e., working from home) brought about by the COVID-19 pandemic. Moreover, we believed the established “rules” in the literature may not specifically apply to our group or to the current somewhat unique conditions. Although the lab members’ values corresponded to reported themes in the literature, lab members felt that the process of deriving our own themes was an incredibly important step in understanding each other’s values and improving our culture significantly beyond what can be achieved by just reading published guidelines.

The self-assessment methodology was effective in increasing satisfaction scores of our lab culture, even compared to before the pandemic when we were working on-site. It seems likely that other approaches to a self-assessment would also be successful, but there were key elements of our approach that we found strongly contributed to the success of our self-assessment:

1. Communication: giving lab members a forum to openly discuss their values
2. Engagement: ensuring each member had the opportunity to participate and share their thoughts with the option of anonymity
3. Leadership: designating session organizers to guide the process
4. Reassessment: evaluating lab member satisfaction at multiple time points and committing to re-evaluating themes as the lab members, projects, and situations change

In summary, we assessed the culture of our lab using anonymous surveys and five 2-3-hour live group sessions, and subsequently improved lab member satisfaction. We identified 10 themes for our group to embody, identified weak points in our lab culture, and then proposed and enacted tangible solutions to address them. While the themes we developed apply uniquely to our group, the success of this team project suggests that other scientific labs could benefit from similar interactive self-assessments.

## Acknowledgments

We would like to thank our advisor, Dr. Laurence Court, for supporting our efforts towards a healthy lab culture. We would like to acknowledge Deborah Mann, who was the mental health champion of our group and was a motivating factor for this study. We would like to thank Dr. Paula Iaeger for helping with survey design. Finally, a thank you to all lab members for committing to this project and staying engaged throughout the entire process.

