## Appendix for "Improving Lab Culture through Self-Assessment: A Case Study" (from another version of this preprint: version 2, 2021-12-11)

**S1 Appendix**

**Pre-Assessment Survey Questions**

1. Assume that you are interviewing for a lab position. What values are important to you in your search for a healthy lab environment. Please identify at least three lab values
2. Assume that you are interviewing for lab positions. What values are important to you in your search for a healthy lab environment? Please enter AT LEAST 3 values. However, we strongly encourage you to enter as many as you can think of.
3. Please rank the values using drag and drop, where choice [1] is the most important value to you.
4. How well did the Court Lab embody these values pre-pandemic?
5. How well do we currently embody these values?

**Post-Assessment Survey Questions**

1. Assume that you are interviewing for lab positions. Out of our Court Lab themes, which ones are most important to you in your search for a healthy lab environment? Please rank/drag our finalized lab themes (with 1 being the theme most meaningful to you).
   a. Accountability + Motivation/Time Management
   b. Adaptability/Crisis Management
   c. Attitude + Fun
   d. Communication
   e. Diversity and Inclusion + Resilience and Success for Diverse Members

f. Emotional Intelligence
g. Ethics/Honesty + Integrity
h. Teamwork/Collaboration
i. Training (Scientific Education, Professional Development, and Professionalism) + Mentorship/Leadership Training + social media
j. Wellbeing + Psychological Safety + Celebrate Success and Normalize Failure

2. Do you have any comments on how these themes are organized or how they are named?
3. Do you think we missed a value that wasn't encompassed by one of these themes? If yes, please explain.
4. Please rate how well you think our lab is currently embodying the finalized themes.
5. Did you like this assessment of our lab culture?
6. Please explain your answer if you would like.
7. What did you think of the time commitment for this process?
8. Please let us know why the time commitment wasn't fully reasonable.
9. Do you feel like your voice was heard throughout this PIAD process?
10. Please help us understand how we could do a better job of ensuring your voice is heard.
11. Should we have another assessment of our lab culture in the future?
12. How often should we assess our lab culture? - conclusion(?)
13. Do you remember any changes that were suggested or carried out?

14. List as many changes as you can remember here.
15. Have there been any changes that you have particularly benefited from?
16. Please talk about the changes you have benefited from and why?
17. What is the likelihood that we will implement these ideas and sustain these changes?
18. What are some ways we can keep up with our themes and continue to improve? In other words, how do we make our changes stick?
19. Is there anything you would change about this assessment process in the future?
20. If you had to pick, which themes would you be interested in helping write about in the paper? Pick 3. (This won't necessarily be final, but it will help in how we split the groups!)
21. Now rank/drag the themes you would write about (1 = First Choice, 2 = Second Choice, 3 = Third Choice)
22. Please rate how well the Court Lab currently embodies your values you submitted last Survey.
23. Do you feel like your concerns about the lab have been addressed?
24. Please explain why you don't think your concerns have been fully addressed.
